# CTP synthase does not form cytoophidia in *Drosophila* interfollicular stalks

**DOI:** 10.1101/2021.12.02.470862

**Authors:** Zheng Wu, Ji-Long Liu

## Abstract

CTP synthase (CTPS) catalyzes the final step of de novo synthesis of the nucleotide CTP. In 2010, CTPS has been found to form filamentous structures termed cytoophidia in *Drosophila* follicle cells and germline cells. Subsequently, cytoophidia have been reported in many species across three domains of life: bacteria, eukaryotes and archaea. Forming cytoophidia appears to be a highly conserved and ancient property of CTPS. To our surprise, here we find that polar cells and stalk cells, two specialized types of cells composing *Drosophila* interfollicular stalks, do not possess obvious cytoophidia. Moreover, we show that Myc level is low in these two types of cells, supporting the idea that Myc regulates cytoophidium assembly. Treatment with a glutamine analog, 6-diazo-5-oxo-l-norleucine (DON), increases cytoophidium assembly in main follicle cells, but not in polar cells or stalk cells. Our findings provide an interesting paradigm for the in vivo study of cytoophidium assembly and disassembly among different populations of follicle cells.

## INTRODUCTION

CTP synthase (CTPS) catalyzes the rate limiting step of de novo CTP synthesis pathway and was found to form intracellular filaments in a wide range of species ^[1], [2], [3], [4], [5], [6], [7]^. These filaments were named as cytoophidia which means ‘cell snakes’^[3]^. CTPS is essential for DNA and RNA metabolism, while CTP is also involved in phospholipid synthesis^[8]^. Previous studies have shown that cytoophidia exist in fast growing cells^[9,10]^. Cytoophidia can also be found in cancer cells^[11]^. These evidences suggest that the formation of cytoophidium is very likely to be related to cell growth and proliferation.

*Drosophila* has an efficient reproductive system and the ovary of the female fly contains many fast-growing cells. Cytoophidia are abundant in the *Drosophila* ovary, making it a good model to study cytoophidium formation and cell growth. The ovary is composed of ovarioles, which are strings of egg chambers. In an egg chamber, somatic follicle cells surround 16 germline cells. These follicle cells proliferate fast during early oogenesis, in which they undergo mitotic divisions. After stage 6, these cells switch to endocycles to grow larger than those at previous stages^[12]^.

A specialized group of follicle cells called stalk cells form stalks to connect between the egg chambers. Polar cells are another group of specialized follicle cells that locate at the anterior and posterior poles of an egg chamber^[13]^. Stalk cells and polar cells function at early oogenesis to separate egg chambers^[14]^. Unlike other follicle cells in early and mid-oogenesis, Stalk cells and polar cells do not proliferate or grow larger after they are differentiated.

Previous studies have demonstrated that the existence of cytoophidia in main body follicle cells is correlated with the growing status of the cells^[2], [10]^. From early oogenesis to stage 10A, there is one cytoophidium in each main body follicle cell, in which the cytoophidium becomes larger as the cell grows bigger. However, whether cytoophidium distributes in polar cells and stalk cells remains unknown. Here, we report that there is no cytoophidium in polar cells and stalk cells. The Myc level in these two kinds of cells is much lower than the other main body follicle cells. This may explain the absence of cytoophidium and the arrest of growth. 6-diazo-5-oxo-L-norleucine (DON) treatment cannot induce cytoophidium assembly in polar cells and stalk cells, suggesting low CTPS level in these cells. These findings provide more evidence for the regulation of Myc on CTPS and cytoophidium assembly in wild-type conditions. In addition, the discovery of low Myc level in polar cells and stalk cells also provides insights for the regulation of cell fate and cell cycles in these cells.

## RESULTS

### Polar cells do no not possess cytoophidium

Cytoophidia is abundant in *Drosophila* ovaries. Generally, there is one cytoophidium in each main body follicle cell in early and mid-stage egg chambers. Polar cells are specialized follicle cells that do not proliferate after differentiation and are arrested at G2 phase^[15]^. In contrast to the existence of cytoophidia in main body follicle cells, we found no cytoophidium in polar cells (Fig.1).

**Figure 1.**
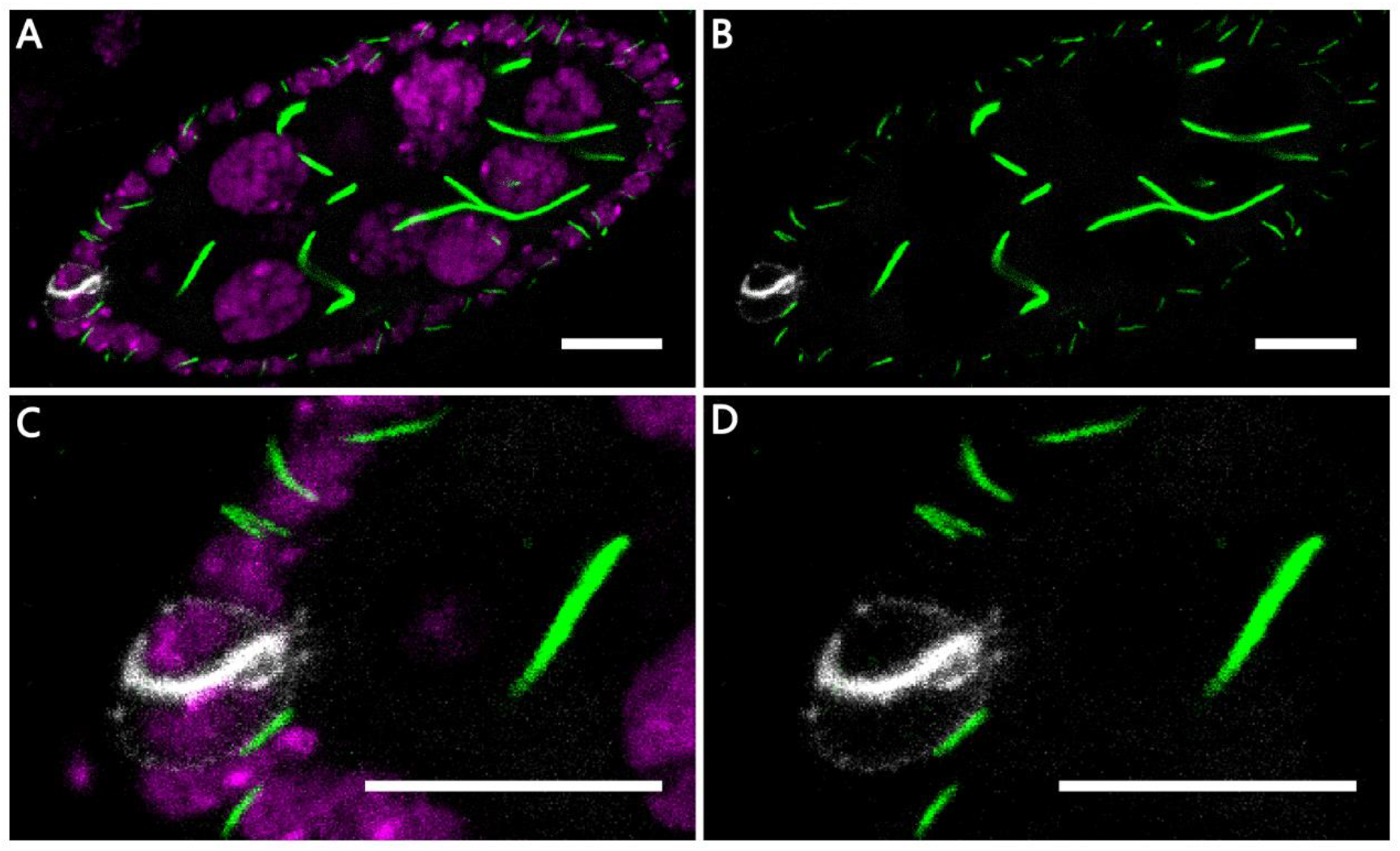
There is no cytoophidium in polar cells. (A,B) DNA (magenta) is stained with Hoechst 33342, cytoophidia (green) are labelled with CTPS antibody. The membrane of polar cells is marked by Fasciclin-3 (white). No cytoophidium can be found in polar cells, while there is one cytoophidium in each of the other main body follicle cells. (C,D) Zoom in of A,B. Scale bars, 10 μm.

We observed cytoophidia in polar cells from other stages and confirmed that no cytoophidium exists in polar cells. Previous studies show that cytoophidia are abundant in fast growing cells^[9], [10]^. Since polar cells are non-proliferating cells, the absence of cytoophidium in polar cells may be related to its growing condition.

### Myc level in polar cells is low

Myc has been found to regulate cytoophidia formation in *Drosophila* follicle cells^[10]^. The staining of *Drosophila* Myc in the ovary revealed that Myc level in polar cells is much lower than the other main body follicle cells in early and mid-oogenesis (Fig.2).

**Figure 2.**
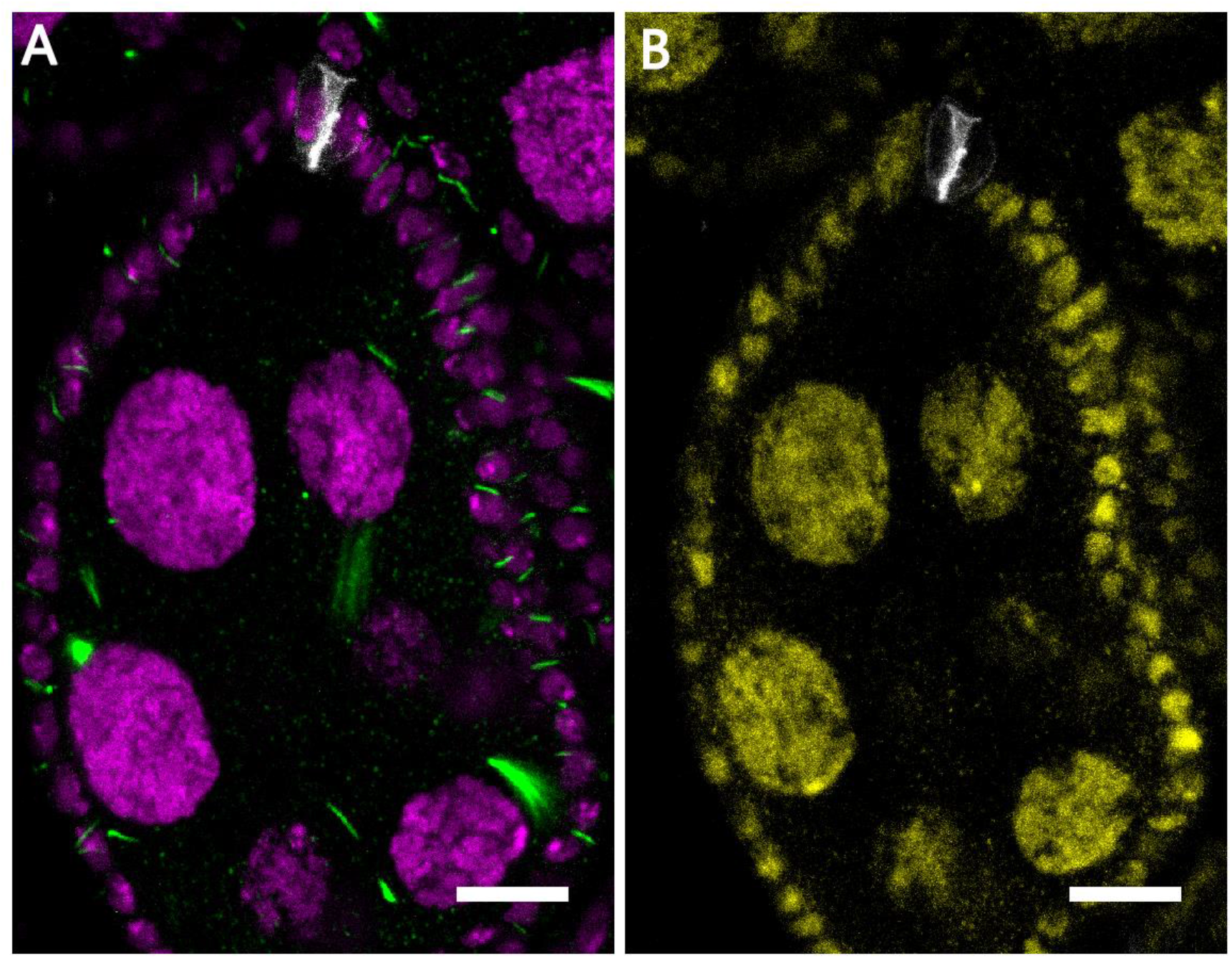
Myc level in polar cells is low. (A) There is no cytoophidium (green) in polar cells (white). DNA is labelled by Hoechst 33342 (Magenta). (B) Myc (yellow) level in polar cells (white) is low, comparing to the other main body follicle cells from the same egg chamber. Scale bars, 10 μm.

The absence of cytoophidium in polar cells is likely to be caused by low Myc level. This finding provides evidence for the correlation of cell proliferation, Myc level and cytoophidium existence.

### Stalk cells do not possess cytoophidium and have low Myc level

Stalk cells are specialized follicle cells that connect the egg chambers. Stalk cells and polar cells are found to be derived from a same group of precursor follicle cells^[16]^. Stalk cells do not proliferate after differentiation and have the function of separating egg chambers^[17]^. We speculate that cytoophidia distribution and Myc level in stalk cells are similar to polar cells.

The staining of CTPS and *Drosophila* Myc confirmed the speculation. There is no cytoophidium in stalk cells and Myc level in stalk cells is very low (Fig.3). This provides another evidence for the regulation of cytoophidia formation by Myc.

**Figure 3.**
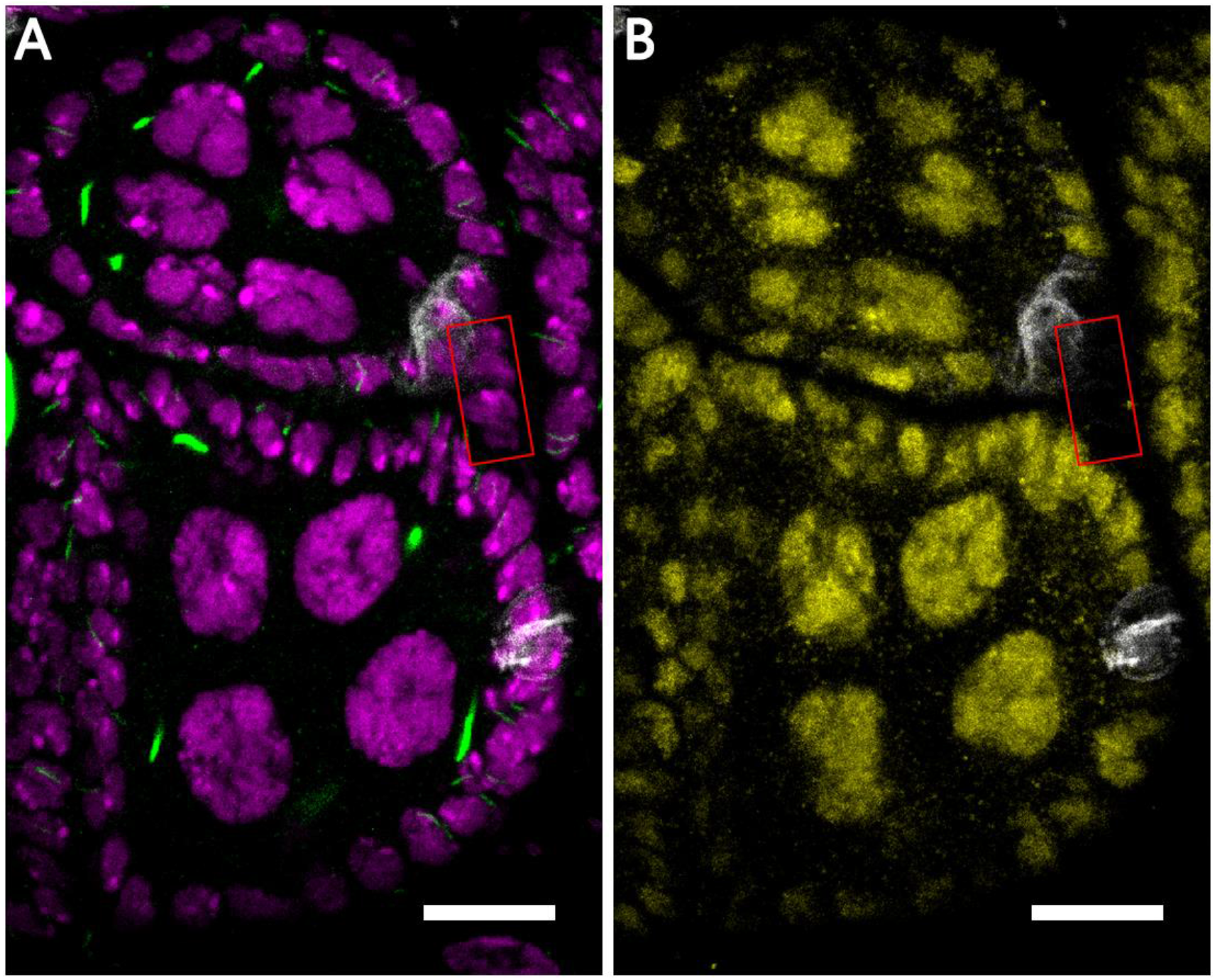
Cytoophidia distribution and Myc level in stalk cells. (A) DNA is labelled by Hoechst 33342 (Magenta). Stalk cells (highlighted by the red box) connect two egg chambers. There is no cytoophidium (green) in stalk cells and polar cells (white). (B) Myc (yellow) level in stalk cells (highlighted by the red box) is much lower than other main body follicle cells. Scale bars, 10 μm.

### DON treatment cannot induce cytoophidium assembly in polar cells and stalk cells

6-diazo-5-oxo-l-norleucine (DON) is a L-glutamine analog. DON can bind to the glutamine amidotransferase domain of CTPS irreversibly and inhibits its activity^[18, 19]^. DON treatment has been found to promote cytoophidium assembly in *Drosophila* ovaries ^[20]^.

We tested whether DON treatment can induce cytoophidium assembly in polar cells or stalk cells. Upon DON treatment, cytoophidia in early and mid-oogenesis have dramatic increasements in length, thickness and number (Fig.4). However, DON treatment cannot induce cytoophidium assembly in polar cells, while CTPS foci sporadically appears in some of the polar cells (Fig.5). Previous research has shown that CTPS foci may represents the early phase of cytoophidium formation^[21]^. The fact that CTP foci can only be found in a few polar cells upon DON treatment indicates that CTPS level in polar cells may be too low to form filaments.

**Figure 4.**
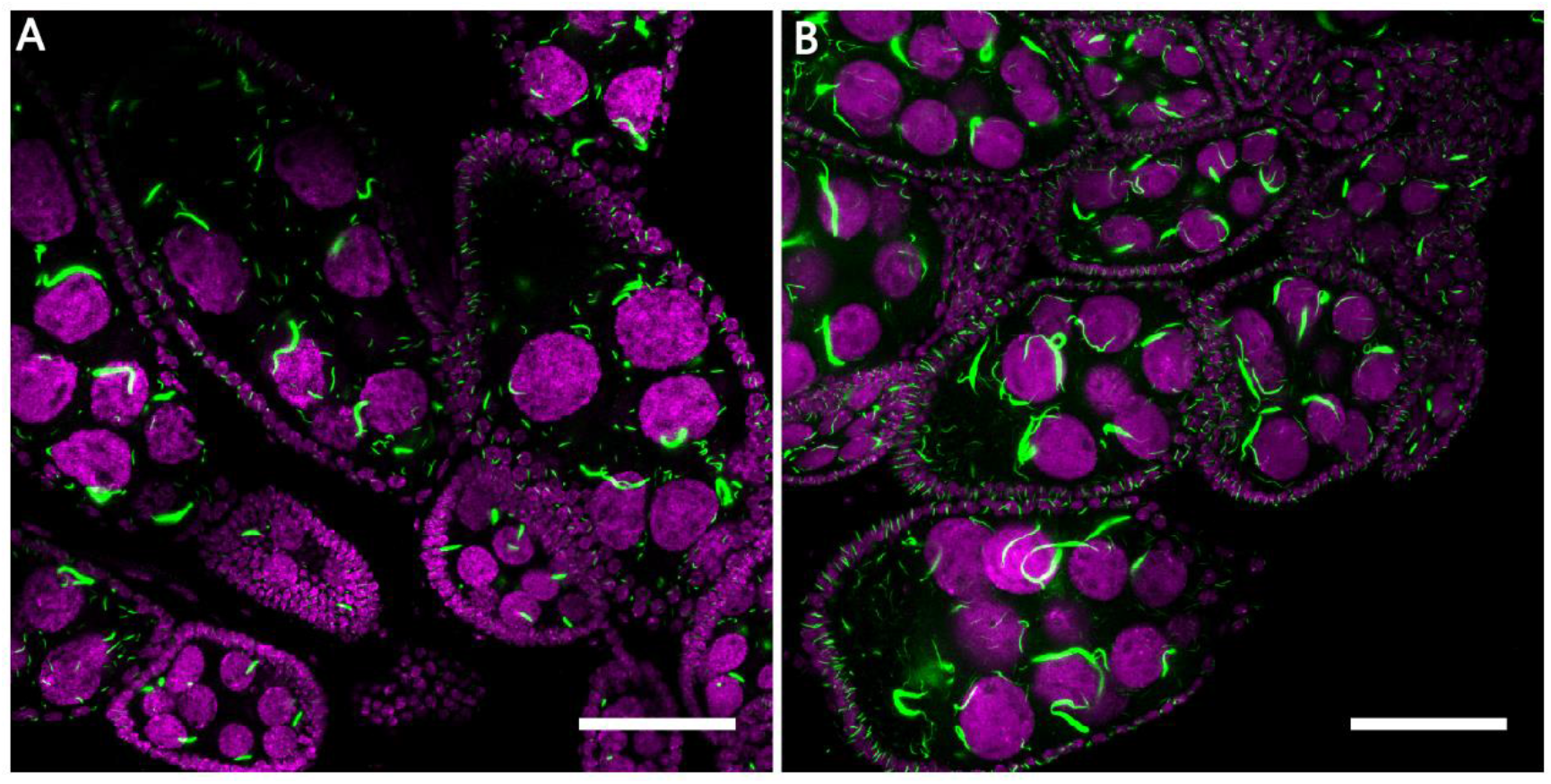
DON treatment promotes cytoophium formation in the early and mid-stage oogenesis. DNA is labelled by Hoechst 33342 (Magenta), cytoophidia (green) are labelled with CTPS antibody. (A) Cytoophidia is abundant in early and mid-stage oogenesis. (B) Upon DON treatment, cytoophidia increase dramatically. Scale bars, 50 μm.

**Figure 5.**
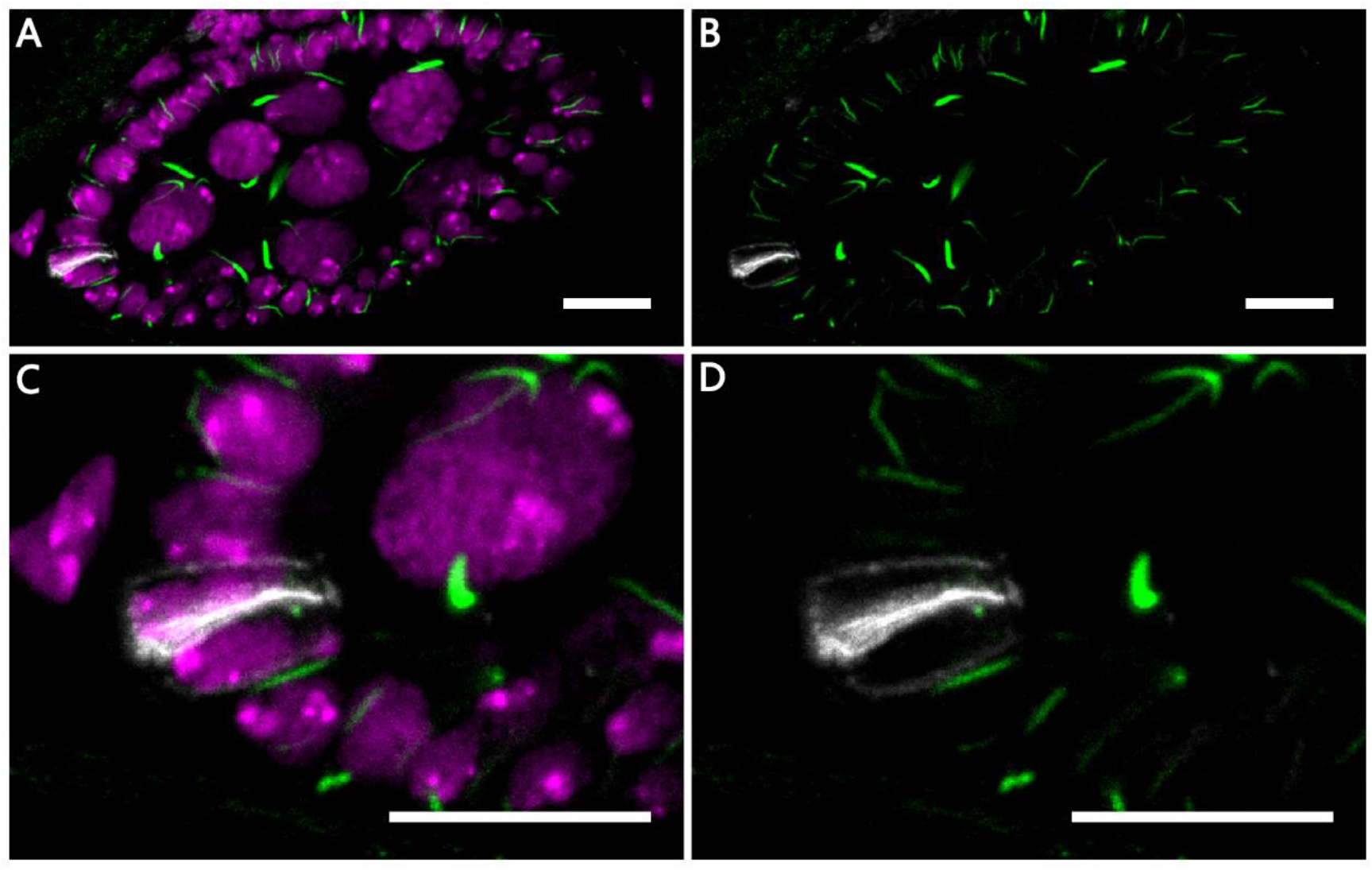
Effect of DON treatment on CTPS in polar cells. (A,B) DNA is labelled by Hoechst 33342 (Magenta). No cytoophidium formed by CTPS (green) can be found in polar cells (white), however CTPS foci (green) can be found in a small portion of the polar cells. (C,D) Zoom in of A,B. Scale bars, 10 μm.

In stalk cells, DON treatment cannot induce cytoophidium or CTPS foci formation (Fig.6). Since previous study shows that DON treatment can induce cytoophidium formation in low CTPS level cells, such as CTPS RNAi knock-down follicle cells^[20]^, the complete absence of cytoophidium or CTPS foci in stalk cells upon DON treatment may indicate extremely low level of CTPS in stalk cells.

Myc has been shown to regulate CTPS expression^[10]^. We have shown that Myc level in polar cells and stalk cells are low, which may result in a much lower CTPS level in these cells than other main body follicle cells. This considerable low level of CTPS then caused the absence of cytoophidium in these cells, even after DON treatment.

**Figure 6.**
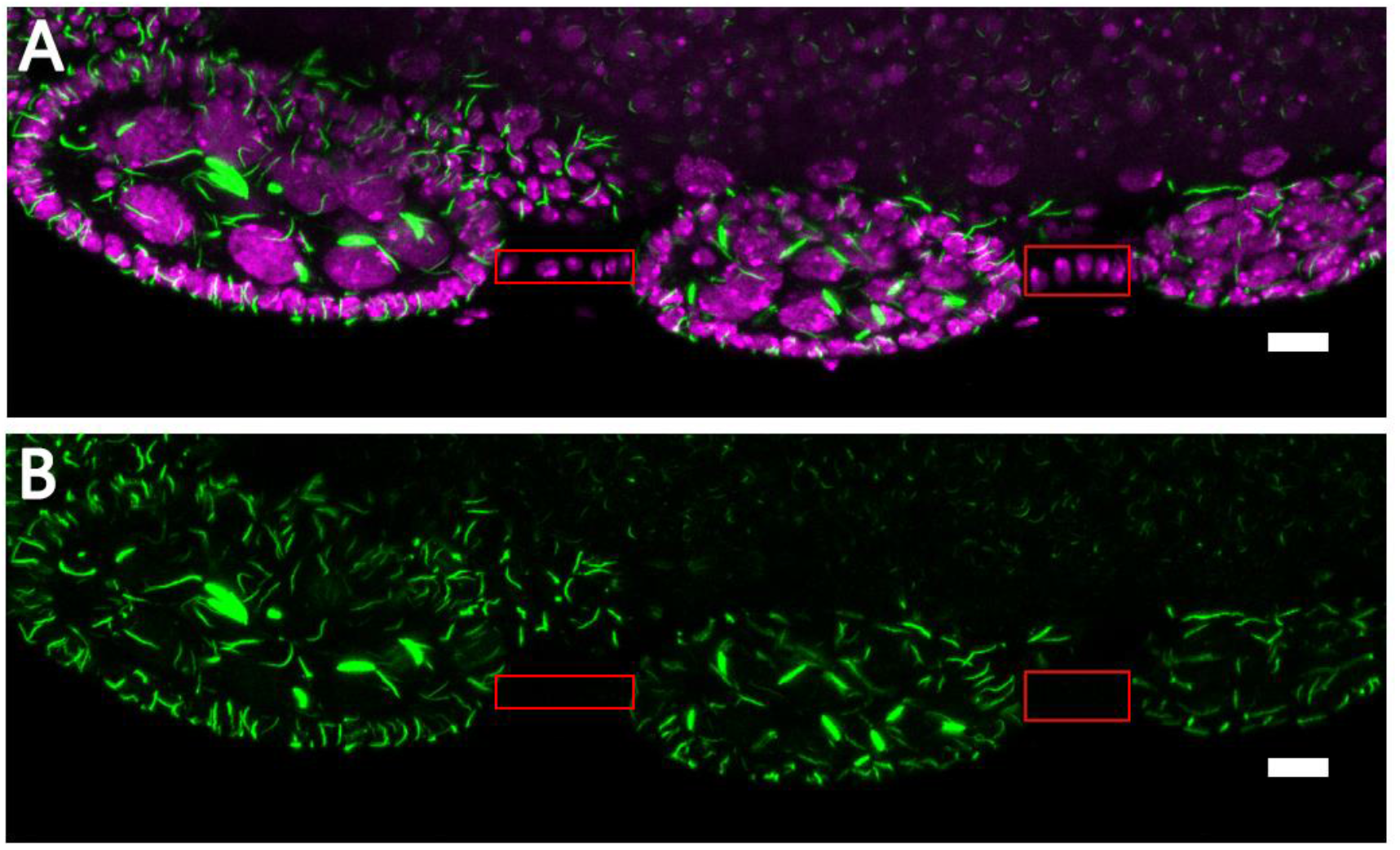
Effect of DON treatment on CTPS in stalk cells. (A,B) DNA is labelled by Hoechst 33342. Stalk cells are highlighted by red boxes. No filaments or foci formed by CTPS (green) can be found in stalk cells. Scale bars, 10 μm.

**Figure 7.**
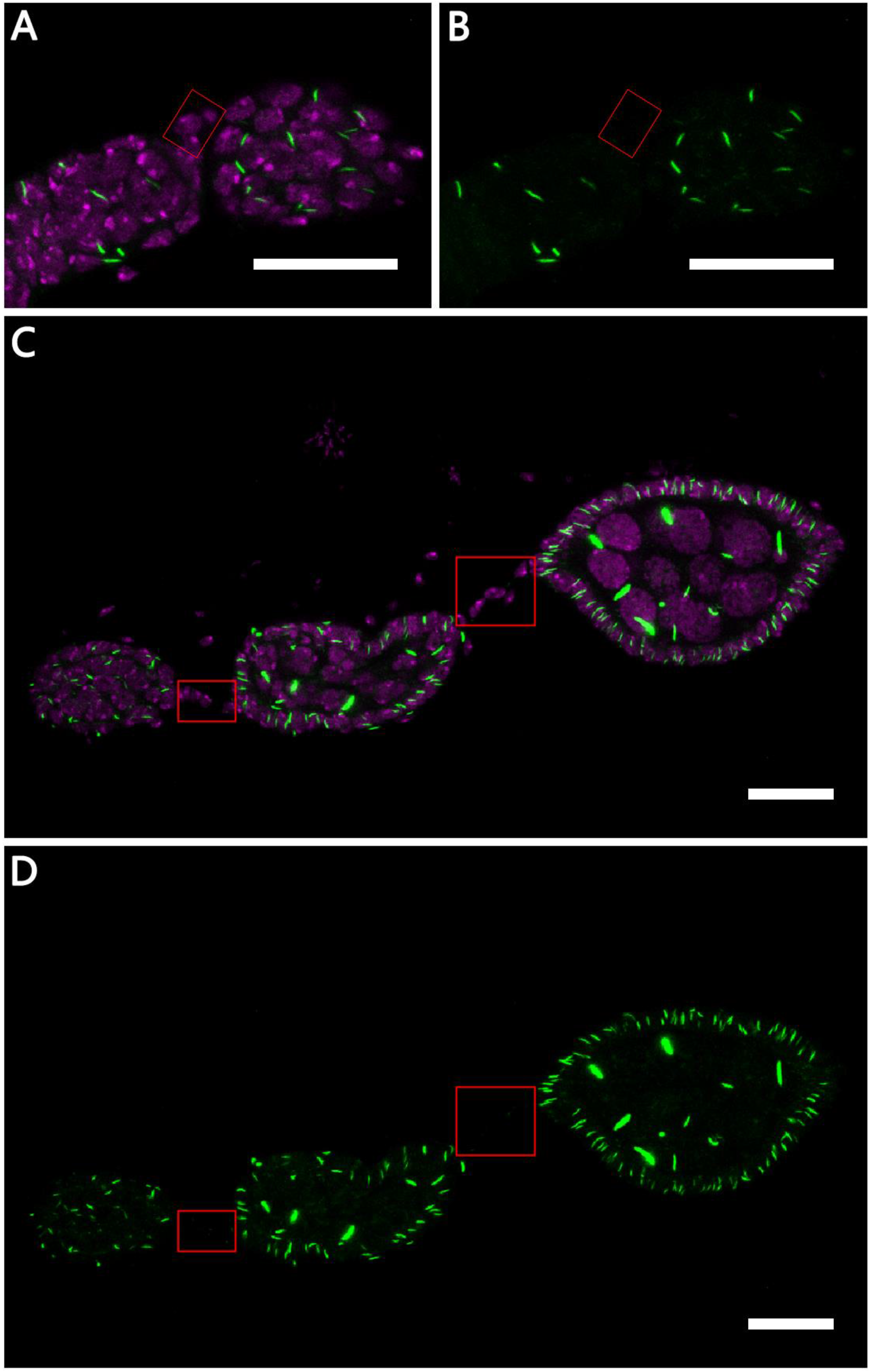
mCherry-tagged CTPS (green) signal confirms no cytoophidium in stalk cells. DNA is labelled by Hoechst 33342 (Magenta). (A,B) Cytoophidium (green) does not distribute in stalk cells (highlighted by the red box) in early oogenesis. (C,D) Stalk cells (highlighted by the red box) connect between egg chambers. No cytoophidium (green) can be found in stalk cells. Scale bars, 20 μm.

## DISCUSSION

Our results show that cytoophidium is absent from polar cells and stalk cells, which are differentiated cells that do not proliferate. A recent study shows that formation of cytoophidium can prolong the half-life of intracellular CTPS molecules^[22]^. These evidences support our speculation that cytoophidium may function as an organelle to store CTPS molecules in fast-growing cells. Follicle cells in the *Drosophila* ovary are all derived from two follicle stem cells, while there is a great difference in the growing status of different kinds of follicle cells. The correlation of cytoophidium existence and fast-growing status suggests that cytoophidium may be considered as an indicator of cell proliferation.

The finding of low Myc level in polar cells and stalk cells is consistent with the previous research of Myc regulation on cell growth and cytoophidium assembly in follicle cells^[10]^. The previous research performed genetic manipulation to up or downregulate the Myc level in main body follicle cells to show the function of Myc. Here we show that Myc is downregulated in two kinds of specialized follicle cells in normal development of the ovary. Besides, low CTPS level in these cells are revealed by DON treatment. Our results provide evidence for the role of Myc in regulating cell growth and CTPS level during follicle cell development. Myc is an important transcriptional factor and it is also well known as an oncogene. CTPS has been found to be upregulated in cancer cells in early research^[23],[24]^. Since cytoophidium has recently been found to form in cancer cells, CTPS could be a potential target for cancer therapy^[11]^. Current we speculate that Myc regulate CTPS expression level and the formation of cytoophidium increases the stability of CTPS. Cancer cells require a large amount of nucleotides for their metabolism and proliferation, so interrupting Myc’s upregulation on CTPS or targeting cytoophidium to decrease CTPS level may be used to restrict tumor growth.

Many researches have been done to study the differentiation of polar cells and stalk cells and several signaling pathways are involved. Hedgehog and Wnt signaling regulates polar cell and stalk cell fate through Eyes absent (Eya) and Castor (Cas)^[25], [26]^. Hippo pathway regulates Notch signaling to maintain polar cell fate, while JAK/STAT signaling are required in the differentiation of stalk cells^[14], [27]^. Our finding of low Myc level in polar cell and stalk cell suggests that the downregulation of Myc could be involved in the signaling pathways mentioned above.

In conclusion, our study shows the correlation of cytoophidium formation, CTPS level, Myc level and cell growth in the development of *Drosophila* ovary. These results further prove the role of Myc in regulating nucleotide metabolism and cytoophidium formation. In addition, the low Myc level and absence of cytoophidium in polar cell and stalk cell provide more insights of cell differentiation in the *Drosophila* ovary for future studies.

## ACKNOWLEDGMENTS

This work was supported by ShanghaiTech University, the UK Medical Research Council (Grant No. MC_UU_12021/3 and MC_U137788471), and National Natural Science Foundation of China (Grant No. 31771490). We thank the Molecular Imaging Core Facility (MICF) at the School of Life Science and Technology, ShanghaiTech University for providing technical support.

## AUTHOR CONTRIBUTIONS

Data curation, Zheng Wu; Formal analysis, Zheng Wu; Funding acquisition, Ji-Long Liu; Investigation, Zheng Wu; Project administration, Ji-Long Liu; Writing – original draft, Zheng Wu; Writing – review & editing, Ji-Long Liu.

## Notes

### Competing Interest Statement

The authors have declared no competing interest.

## REFERENCES

[1] Lieberman, I.. (1956). Enzymatic amination of uridine triphosphate to cytidine triphosphate. Journal of Biological Chemistry, 222(2), 765.

[2] Liu, J. L.. (2010). Intracellular compartmentation of ctp synthase in drosophila. Journal of Genetics and Genomics, 37(5), 281–296.

[3] Ingerson-Mahar, M., Briegel, A., Werner, J. N., Jensen, G. J., & Gitai, Z.. (2010). The metabolic enzyme ctp synthase forms cytoskeletal filaments. NATURE CELL BIOLOGY, 12(8), 739–746.

[4] Noree, C., Sato, B. K., Broyer, R. M., & Wilhelm, J. E. (2010). Identification of novel filament-forming proteins in Saccharomyces cerevisiae and Drosophila melanogaster. The Journal of cell biology, 190(4), 541–551.

[5] Carcamo, W. C., Minoru, S., Hideko, K., Naohiro, T., Takashi, H., & Chan, J. Y. F., et al. (2011). Induction of cytoplasmic rods and rings structures by inhibition of the ctp and gtp synthetic pathway in mammalian cells. PLoS ONE, 6(12), e29690..

[6] Chen, K., Zhang, J., ?mür Yilmaz Tastan, Deussen, Z. A., Siswick, Y. Y., & Liu, J. L.. (2011). Glutamine analogs promote cytoophidium assembly in human and drosophila cells. Journal of Genetics and Genomics, 38(9), 391–402.

[7] Zhang, J., Hulme, L., & Liu, J. L.. (2014). Asymmetric inheritance of cytoophidia in schizosaccharomyces pombe. Biology Open, 3(11), 1092–1097.

[8] Chang Y-F, Carman GM. CTP synthetase and its role in phospholipid synthesis in the yeast Saccharomyces cerevisiae. Progress in lipid research. 2008;47(5):333–339.

[9] Tastan OY., & Liu, J. L.. (2015). Ctp synthase is required for optic lobe homeostasis in drosophila. Journal of Genetics and Genomics, 42(5), 261–274.

[10] Aughey, G. N., Grice, S. J., & Liu, J. L. (2016). The interplay between myc and ctp synthase in drosophila. Plos Genetics, 12(2), e1005867.

[11] Chang, C. C., Jeng, Y. M., Peng, M., Keppeke, G. D., Sung, L. Y., & Liu, J. L. (2017). Ctp synthase forms the cytoophidium in human hepatocellular carcinoma. Experimental Cell Research, 361(2).

[12] Deng, W. M., Althauser, C., & Ruohola-Baker, H.. (2002). Notch-delta signaling induces a transition from mitotic cell cycle to endocycle in drosophila follicle cells. Development, 128(23), 4737–4746.

[13] Margolis, J., & Spradling, A. C.. (1995). Margolis, j. & spradling, a. identification and behavior of epithelial stem cells in the drosophila ovary. development 121, 3797-3807. Development, 121(11), 3797–3807.

[14] Assa-Kunik, E., Torres, I. L., Schejter, E. D., Johnston, D. S., & Shilo, B. Z.. (2007). Drosophila follicle cells are patterned by multiple levels of notch signaling and antagonism between the notch and jak/stat pathways. Development, 134(6), 1161–1169.

[15] Shyu, L. F., Sun, J., Chung, H. M., Huang, Y. C., & Deng, W. M.. (2009). Notch signaling and developmental cell-cycle arrest in drosophila polar follicle cells. Molecular Biology of the Cell, 20(24), 5064–5073.

[16] Tworoger, M., Larkin, M. K., Bryant, Z., & Ruohola-Baker, H. (1999). Mosaic analysis in the drosophila ovary reveals a common hedgehog-inducible precursor stage for stalk and polar cells. Genetics, 151(2), 739–748.

[17] Wu, X., Tanwar, P. S., & Raftery, L. A. (2008). Drosophila follicle cells: morphogenesis in an eggshell. Seminars in cell & developmental biology, 19(3), 271–282.

[18] Levitzki, A., Stallcup, W. B., & Koshland, D. E.. (1971). Half-of-the-sites reactivity and conformational states of cytidine triphosphate synthetase. Biochemistry, 10(18), 3371–3378.

[19] Levitzki, A., & Koshland, D. E.. (1971). Cytidine triphosphate synthetase. covalent intermediates and mechanisms of action. Biochemistry, 10(18), 3365–3371.

[20] Chen, K., Zhang, J., ?mür Yilmaz Tastan, Deussen, Z. A., Siswick, Y. Y., & Liu, J. L.. (2011). Glutamine analogs promote cytoophidium assembly in human and drosophila cells. Journal of Genetics and Genomics, 38(9), 391–402.

[21] Gou, K. M., Chang, C. C., Shen, Q. J., Sung, L. Y., & Liu, J. L.. (2014). Ctp synthase forms cytoophidia in the cytoplasm and nucleus. Experimental Cell Research, 323(1), 242–253.

[22] Sun Z, Liu JL. Forming cytoophidia prolongs the half-life of CTP synthase. (2019). Cell Discov, 5: 32.

[23] Williams JC, Kizaki H, Weber G, Morris HP (1978) Increased CTP synthetase activity in cancer cells. Nature 271: 71–73.

[24] van den Berg AA, van Lenthe H, Busch S, de Korte D, Roos D, et al. (1993) Evidence for transformation-related increase in CTP synthetase activity in situ in human lymphoblastic leukemia. Eur J Biochem 216: 161–167.

[25] Chang, Y. C., Jang, C. C., Lin, C. H., & Montell, D. J.. (2013). Castor is required for hedgehog-dependent cell-fate specification and follicle stem cell maintenance in drosophila oogenesis. Proceedings of the National Academy of Sciences, 110(19), E1734–E1742.

[26] Dai, W., Peterson, A., Kenney, T., Burrous, H., & Montell, D. J.. (2017). Quantitative microscopy of the drosophila ovary shows multiple niche signals specify progenitor cell fate. Nature Communications, 8(1), 1244.

[27] Chen, H. J., Wang, C. M., Wang, T. W., Liaw, G. J., Hsu, T. H., & Lin, T. H., et al. (2011). The hippo pathway controls polar cell fate through notch signaling during drosophila oogenesis. Developmental Biology, 357(2), 370–379.

